# The Effect of Self-Loading on the Mechano-Stability and Stalk Lodging Resistance of Plant Stems

**DOI:** 10.1101/2020.03.21.001727

**Authors:** Christopher J Stubbs, Yusuf Oduntan, Tyrone Keep, Scott D Noble, Daniel J. Robertson

## Abstract

**Background:** Stalk lodging (breaking of agricultural plant stalks prior to harvest) is a multi-billion dollar a year problem. Stalk lodging occurs when bending moments induced by a combination of external loading (e.g. wind) and self-loading (e.g. the plant’s own weight) exceed the bending strength of plant stems. Previous biomechanical plant stem models have investigated both external loading and self-loading of plants, but have evaluated them as separate and independent phenomena. However, these two types of loading are highly interconnected and mutually dependent. The purpose of this paper is twofold: (1) to investigate the combined effect of external loads and plant weight on the displacement and stress state of plant stems / stalks, and (2) to provide a generalized framework for accounting for self-weight during mechanical phenotyping experiments used to predict stalk lodging resistance.

**Results:** A method of properly accounting for the interconnected relationship between self-loading and external loading of plants stems is presented. The interconnected set of equations are used to produce user-friendly applications by presenting (1) simplified self-loading correction factors for a number of common external loading configurations of plants, and (2) a generalized Microsoft Excel framework that calculates the influence of self-loading on crop stems. The effect of self-loading on the structural integrity of wheat is examined in detail. A survey of several other plants is conducted and the influence of self-loading on their structural integrity is also presented.

**Conclusions:** The self-loading of plants plays a potentially critical role on the structural integrity of plant stems. Equations and tools provided herein enable researchers to account for the plant’s weight when investigating the flexural rigidity and bending strength of plant stems.

## Introduction

Yield losses due to stalk lodging (breakage of crop stems or stalks prior to harvest) are estimated to range from 5-20% annually [1,2]. Despite a growing body of literature surrounding the topic of stalk lodging in wheat, barley, oats, and maize [3–7], a detailed study on the interconnected relationship between external loading (e.g. wind) and self-loading (e.g. plant weight) on stalk bending strength has not been reported. Previous biomechanical plant stem models have examined the influence of morphology, material, and weight on stem failure, while others have separately analyzed the effects of externally induced bending forces (e.g., wind) on stem failure [3,4,8–15]. However, the effects of self-weight and external loads on stem failure are inextricably connected. The bending moment induced from self-weight is a function of the distance between the plant’s base and its center of gravity. As external loads displace the center of gravity away from the base of the stem, the bending moment induced from self-weight increases. Although previous studies have independently looked at external loads [4,5,16] and self-weight loading [3], the authors are not aware of previous investigations into the interplay between these factors.

Calculating the total applied bending moment (M_TOTAL_), resulting from the combination of external bending loads (M_ext_) and bending loads induced from self-weight (M_int_) is necessary to understand stem strength and stalk lodging resistance. Stalk lodging resistance is commonly estimated through the use of mechanical phenotyping equipment [e.g.,[6]]. Such equipment typically applies an external load to plant stems and measures plant deflection. The externally applied moment (M_ext_) and plant displacement (*δ*) data are then analyzed to determine the flexural stiffness of the stem (EI) or the bending strength of the stem. For example see [17]:

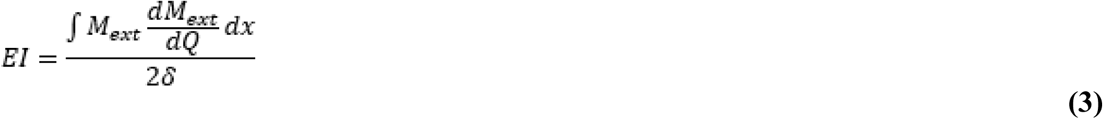

However, the effect of self weight on plant deflection is ignored or assumed to be negligible in these analyses. In some instances researchers have removed leaf blades, grain heads, and all plant biomass above the grain prior to mechanical testing to limit the effect of self-weighting. However, it is unclear if such measures are adequate. To more accurately characterize flexural stiffness and stalk bending strength the contribution of self weight to plant deflection and to the induced bending moment M_TOTAL_ needs to be calculated.

The purpose of this paper is therefore twofold: (1) to investigate the combined effect of external loads and plant weight on the displacement and stress state of plant stems, and (2) to provide a generalized framework for accounting for self-weight during mechanical phenotyping experiments used to predict stalk lodging resistance.

## Methods

### Box 1

**Glossary of Terms**

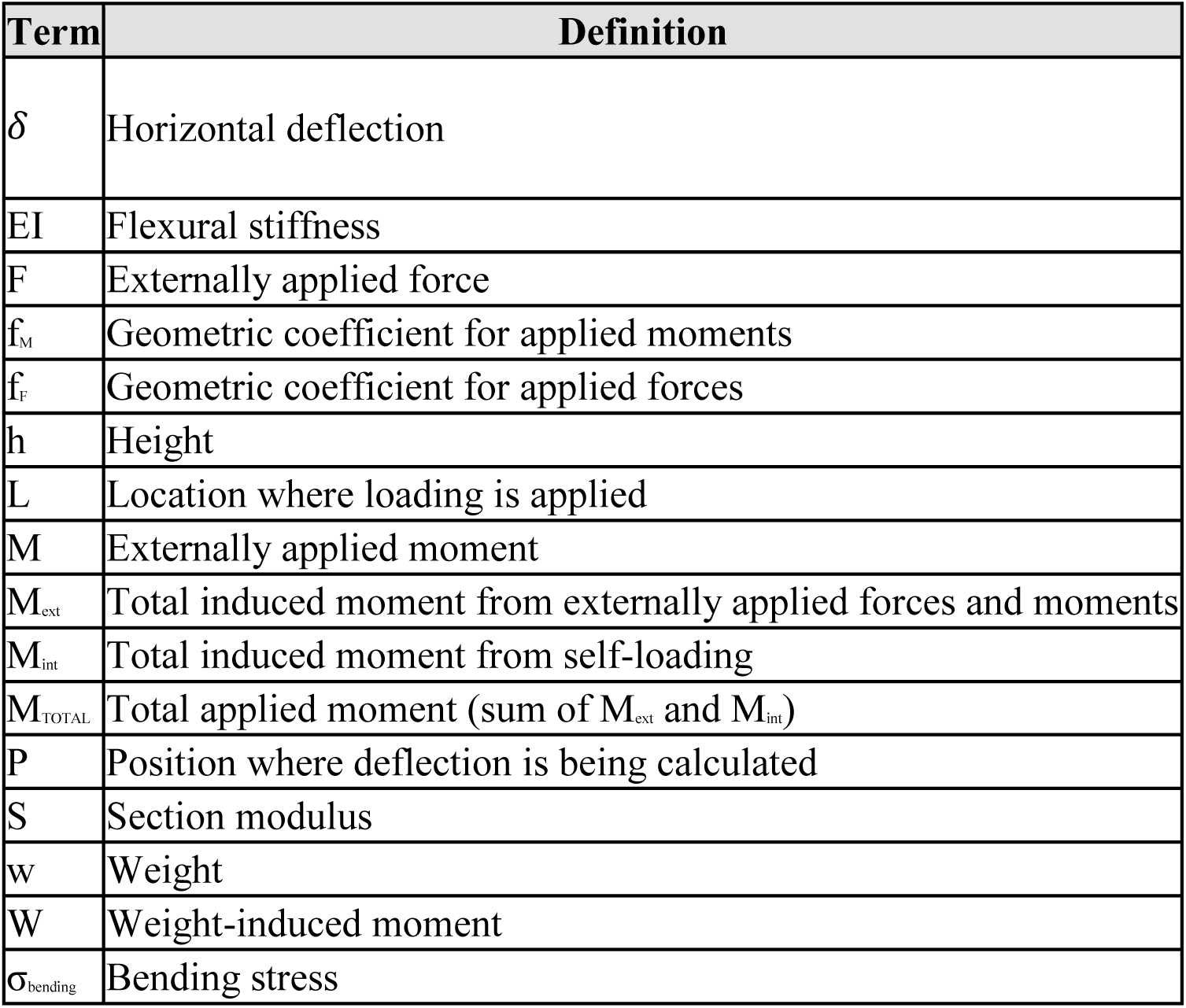

### Derivation of Closed Form Solution

To determine the contribution of self weight to the mechano-stability of plant stems, we must first derive a closed form solution for the internal bending moment of the stem (M_TOTAL_). Figure 1 depicts the free body diagram of a plant stem with an arbitrary loading applied at two locations. This stem depicts two weights (w) (e.g. stem weight, grain weight), as well as two externally applied loads (F) and two externally applied moments (M).

**Figure 1:**
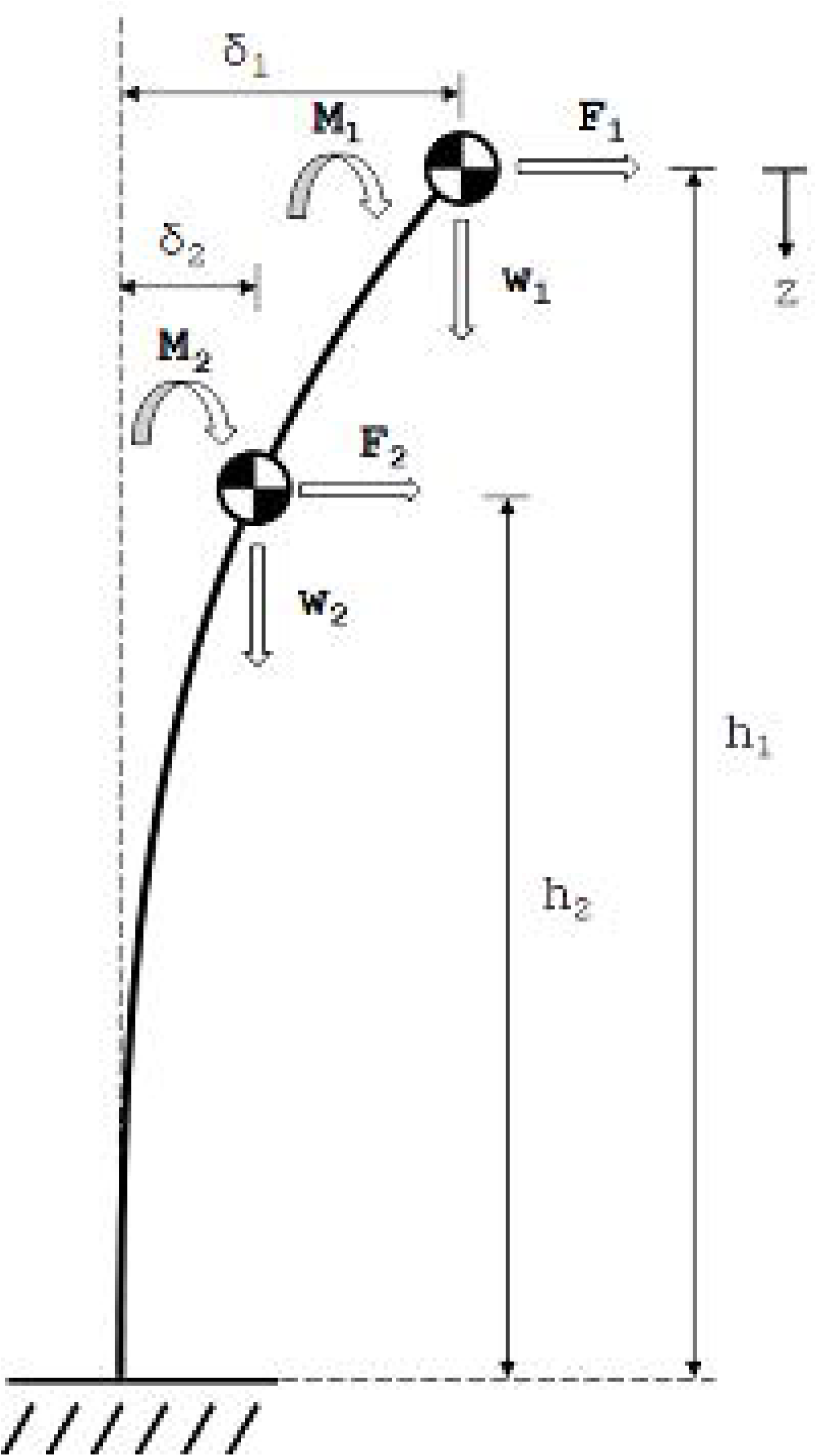
The loading diagram of a deflected stem, showing two loading locations with all three types of loading (an applied force, an applied moment, and a weight).

As the stem deflects, the moments induced from self-weights will increase as a function of the deflection of the stem. For the weight (w) at each location, we can calculate the induced moment from self-weight (W) as the product of the weight and the weight’s offset (i.e., the deflection of the stem at the location of the weight (*δ*)). Thus for the two locations shown in Figure 1, we have:

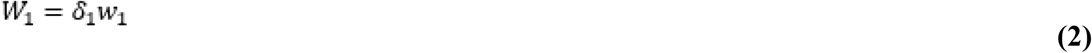

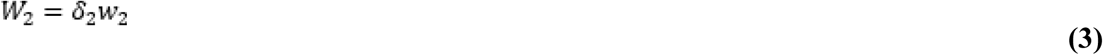

It should be noted that Equations 1, 2 and 6 assume that the maximum moment induced by self-loading is applied to the entire length of the stem. For more detail, see the Limitations section.

However, the offsets (*δ*_1_ and *δ*_2_) in equations 2 and 3 are not known and are a function of the externally applied moments and forces. Using engineering theory for beam deflection [18] and the theory of superposition of loading [17], we can calculate the deflection of the stem at height h_1_ (i.e., location 1) as a function of the applied forces, applied moments, and weight-induced moments. Equation 4 shows this calculation, where the first row of equation 4 concerns loads, moments and weights at location 1 (i.e., at height h_1_) and the second row of equation 4 concerns forces, moments and weights at location 2 (i.e., at height h_2_).

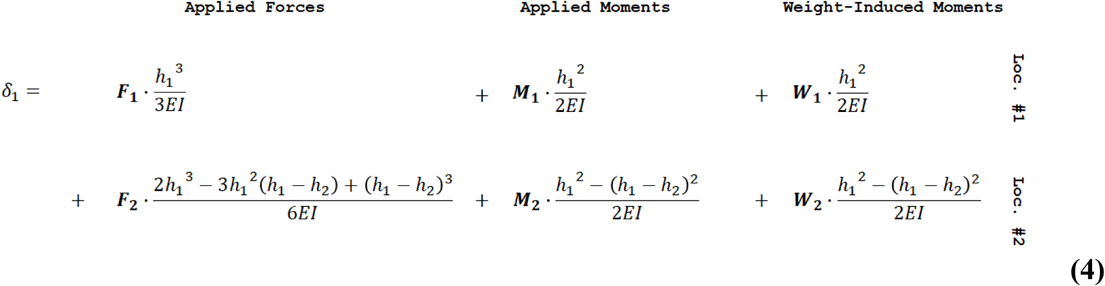

Similarly, we can write the deflection of the stem at h_2_ as:

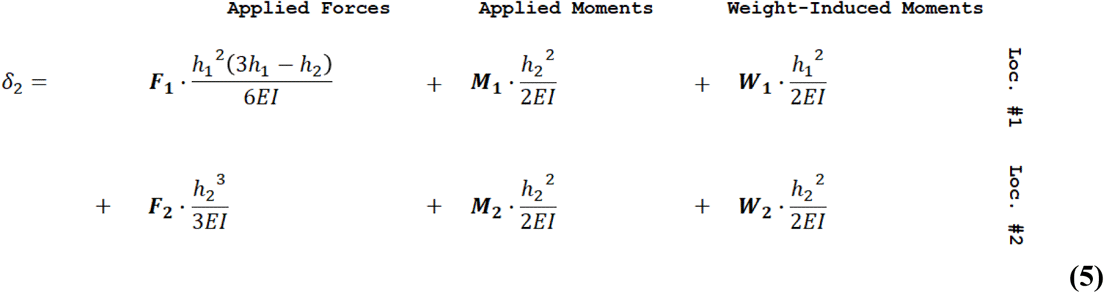

Thus we have four linearly independent equations (Equations 2 through 5) allowing us to solve for four unknown values (W_1_, W_2_, *δ*_1_, *δ*_2_).

Furthermore, Equations 2 through 5 can be generalized to account for any number of locations (n) along the length of the stalk. First, for any loading location L, at a height h_L_ along the stalk, deflected by *δ*_L_, Equations 2 and 3 can be generalized as:

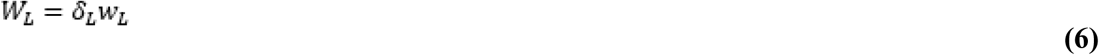

Next, Equations 4 and 5 can be generalized by noting that each force, moment or weight (F, M, or W, shown in bold in Equations 4 and 5) is multiplied by a geometric coefficient. The geometric coefficient for each term is a function of the height where the deflection is measured and the height at which the loading is applied. This geometric coefficient can be denoted as either *f*_F_ (for forces) or *f*_M_ (for applied moments or weight-induced moments). As such, for any location P at a height of h_P_, the deflection *δ*_P_ is calculated by summing the product of each load, moment or weight (F, M, or W) and its corresponding geometric coefficient (*f*_F_ or *f*_M_) at every loading location (from L=1 to L=n). Note that this geometric coefficient assumes a constant flexural stiffness (EI), as discussed in the Limitations section. Thus the generalized form of Equations 4 and 5 can be written as:

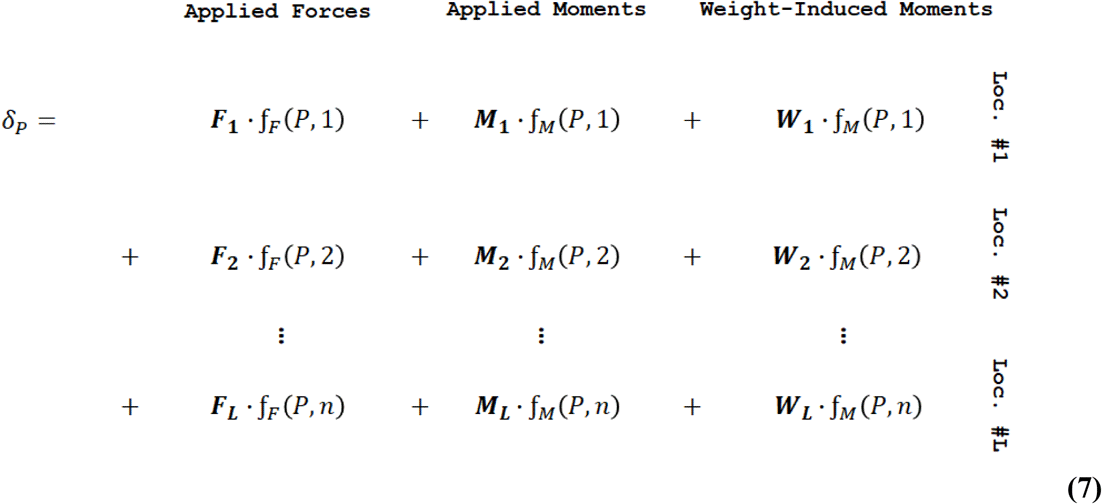

Equation 7 can now be consolidated into a fully generalized form of:

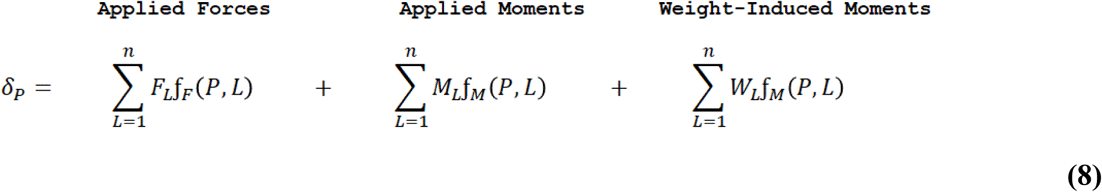

Where the geometric coefficients for the forces and moments are defined as [18]:

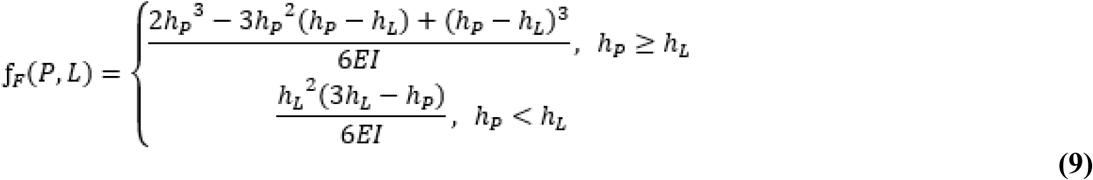

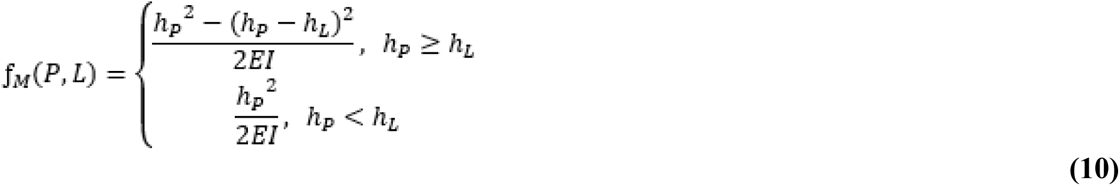

Equations 6 through 9 can also be put into a generalized matrix form. From Equations 6 and 8 we see that for any number of weights at any number of locations (n), we will have *2n* unknown values (*δ*_1_, *δ*_2_, … *δ*_n_, W_1_, W_2_,.. W_n_), and *2n* linearly independent equations. By rearranging these equations and converting them to matrix notation we can write:

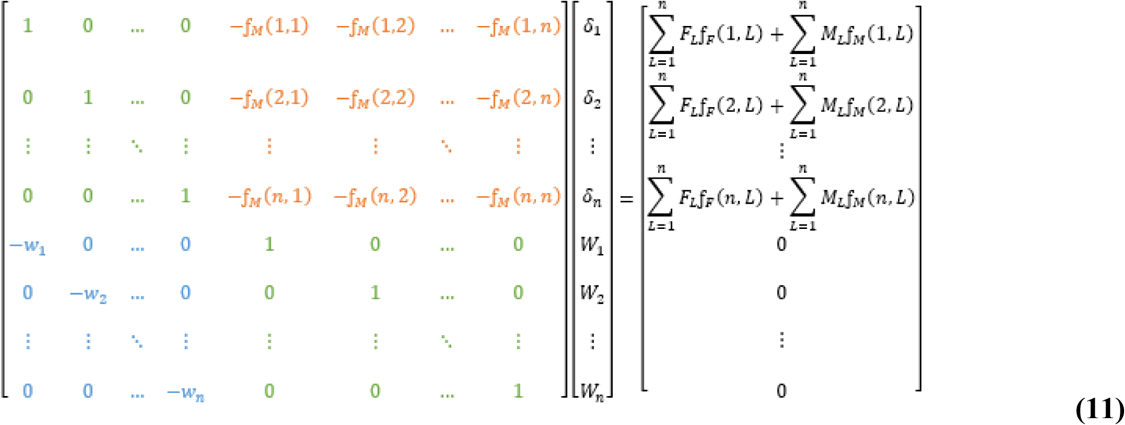

Where the first matrix in the equation is a square matrix of size *2n* x *2n*, and the second and third matrices in the equation are column matrices of size *2n* x *1*. Within the square matrix, the top left and bottom right *n* x *n* submatrices (shown in green text) are identity matrices, the bottom left *n* x *n* submatrix (shown in blue text) is a diagonal matrix of the negative weights (-w), and the top right *n* x *n* submatrix (shown in orange text) is the negative geometric coefficients of the weight-induced moments, as calculated by Equation 10. We can then solve this matrix equation to calculate the deflections and weight-induced moments:

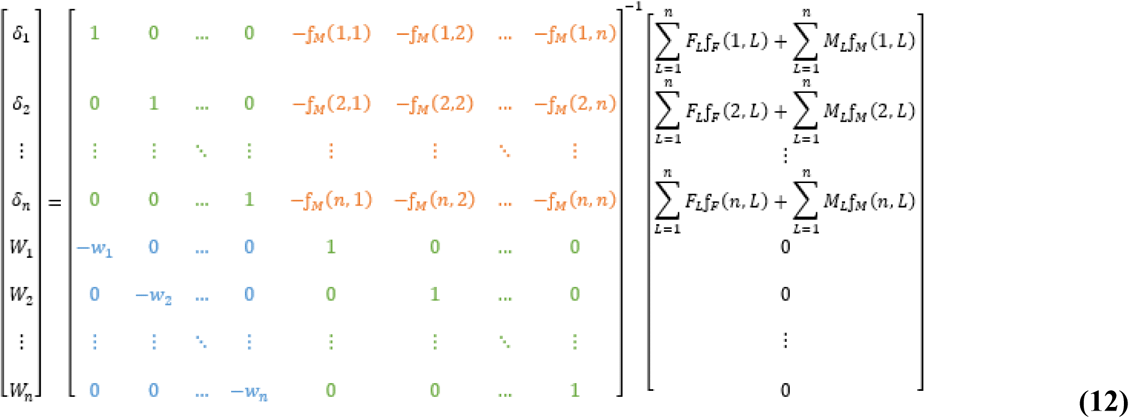

We can now look at the total moment (M_TOTAL_) of any cross-section along the length of the stem. In particular, M_TOTAL_ can be written as a function of h_P_ and h_L_, by considering all of the loads that are applied to the stem above the cross-section of interest (i.e, for h_L_ ≥ h_P_),

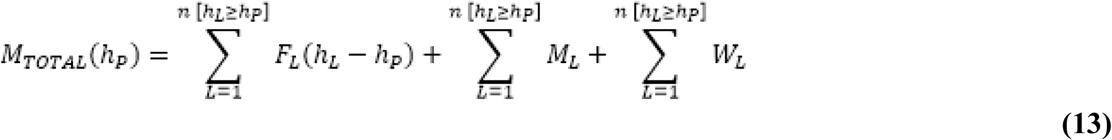

Finally, we can write the bending stress of the stem in terms of the internal moment and the section modulus of the cross-section (S(h_P_)):

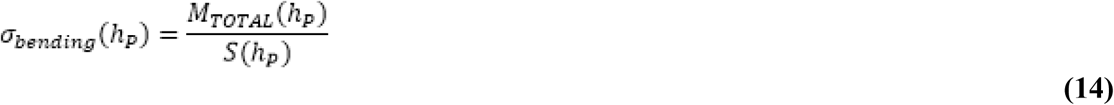

### Finite Element Modeling and Data Triangulation

As a form of data triangulation [18] to confirm the closed form solution presented above, a series of 768 non-linear finite element models of plant stems were developed. The stems were modeled as 2-noded linear beam elements, fixed at their base. The models were developed in Abaqus/CAE 2019 [19,20] and analyzed in Abaqus/Standard 2019 using a direct, full Newton solver [19,20]. Model development and post-processing were automated through a series of custom Python scripts. Stems were modeled with a weight at height h_1_, applied force at height h_2_, and moment at height h_3_. A full parametric sweep of elastic moduli, moment of inertia, heights, moments, weights, and forces were performed. Table 1 describes the parameter space for the models. Values of input parameters to the models were based on previous studies of plant stem material properties [21, 22]. The applied moment (Mtotal) and the deflection of each finite element model was then compared to the deflection and applied moment calculated using the closed form solution methods presented above.

**Table 1:**
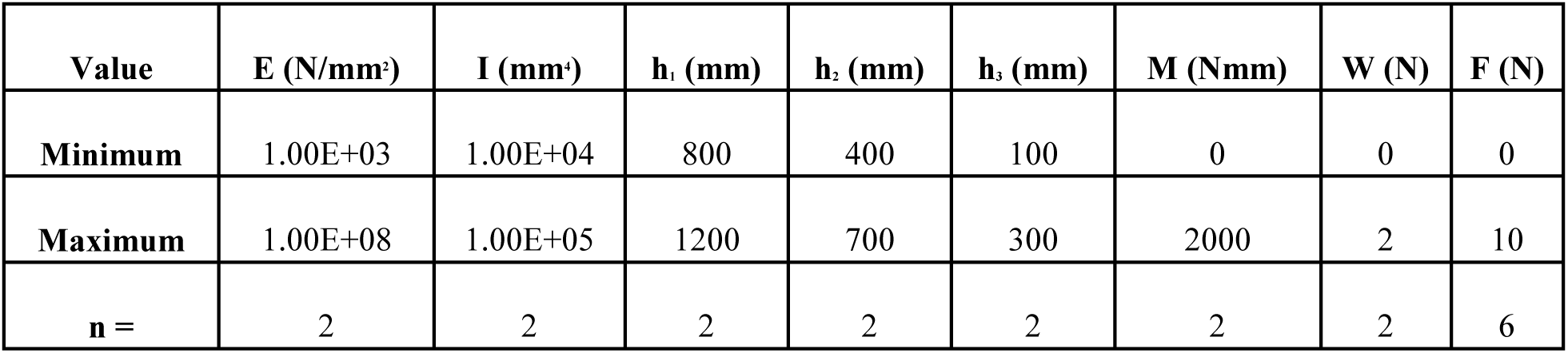
The parameter space of the finite element analyses. Models were developed for the minimum and maximum values (n=2), and evaluated every 2N of applied load from 0N to 10N (n=6). A total of 768 finite element models were evaluated.

### Analyzing the Effect of Self-Loading on Wheat

The closed form solution method was applied to vertically-partitioned wheat biomass data to determine the effects of self loading in wheat. Biomass data was collected from a commercially available wheat (*Triticum aestivum*) variety during the 2018 growing season in Saskatoon, Saskatchewan. As depicted in Figure 2 the biomass of wheat stems, leaves, and spikes were sampled and weighed every 10 cm along the length of the plants. Planting density was approximately 1.3 million plants per hectare with a 30.5 cm (12”) row spacing.

**Figure 2:**
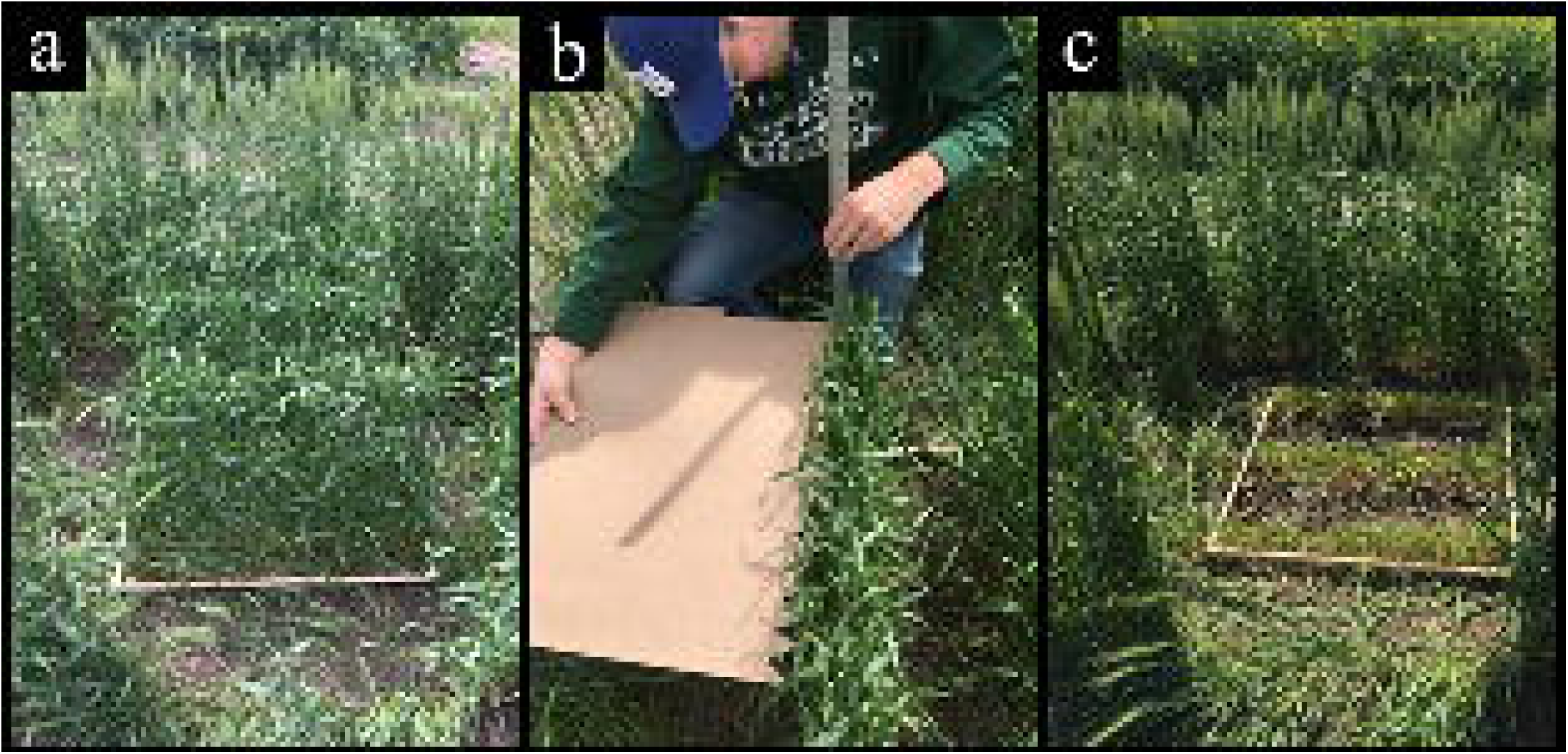
To obtain biomass samples a square sample area (a) was chosen and then cleared of surrounding plants. The sampling boundary height was measured (b) and then the biomass above this layer was cut and bagged. This was continued in 10 cm layers until there was no biomass left (c).

Biomass data were gathered from a 68 cm x 68 cm square of wheat in the center of a 122 cm wide plot. The 68 cm x 68 cm square contained an average of 366 stems. A total of five samples of biomass data were taken periodically from July 27 to August 29, 2018. The same plot was used for all sampling dates with enough space left between samples to have undisturbed wheat in each subsequent sample. A square guide was placed over the middle rows of the plot to indicate the sampling area and any plants outside of the guide were then removed. Biomass was harvested in 10 cm layers measured from the ground with the highest layer collected first (topmost layer varied in size depending on total plant height). All plant matter from a single layer was harvested, weighed in the field to obtain wet-basis biomass, and bagged to be dried later. The samples were oven dried at 65oC for a minimum of 48 hours to obtain the dry-basis biomass.

## Results

### Comparison of Finite Element and Closed Form Solutions

As a form of data triangulation finite element models of plant stems were compared to the closed form solution presented in the methods section. The finite element models were found to be in good agreement with the closed form solutions (see Figure 3). In particular, the median error between the 768 finite element models and the corresponding closed form solutions was found to be 0.126% for displacement at the top of the specimen, and 0.0003% for the total moment at the base of the specimen. These data imply that for the ranges evaluated, the closed form solution is providing accurate results. It should also be noted that because the displacement is based upon the integral of the moment [17], it is expected that greater error exists in the displacement than the induced moment.

**Figure 3:**
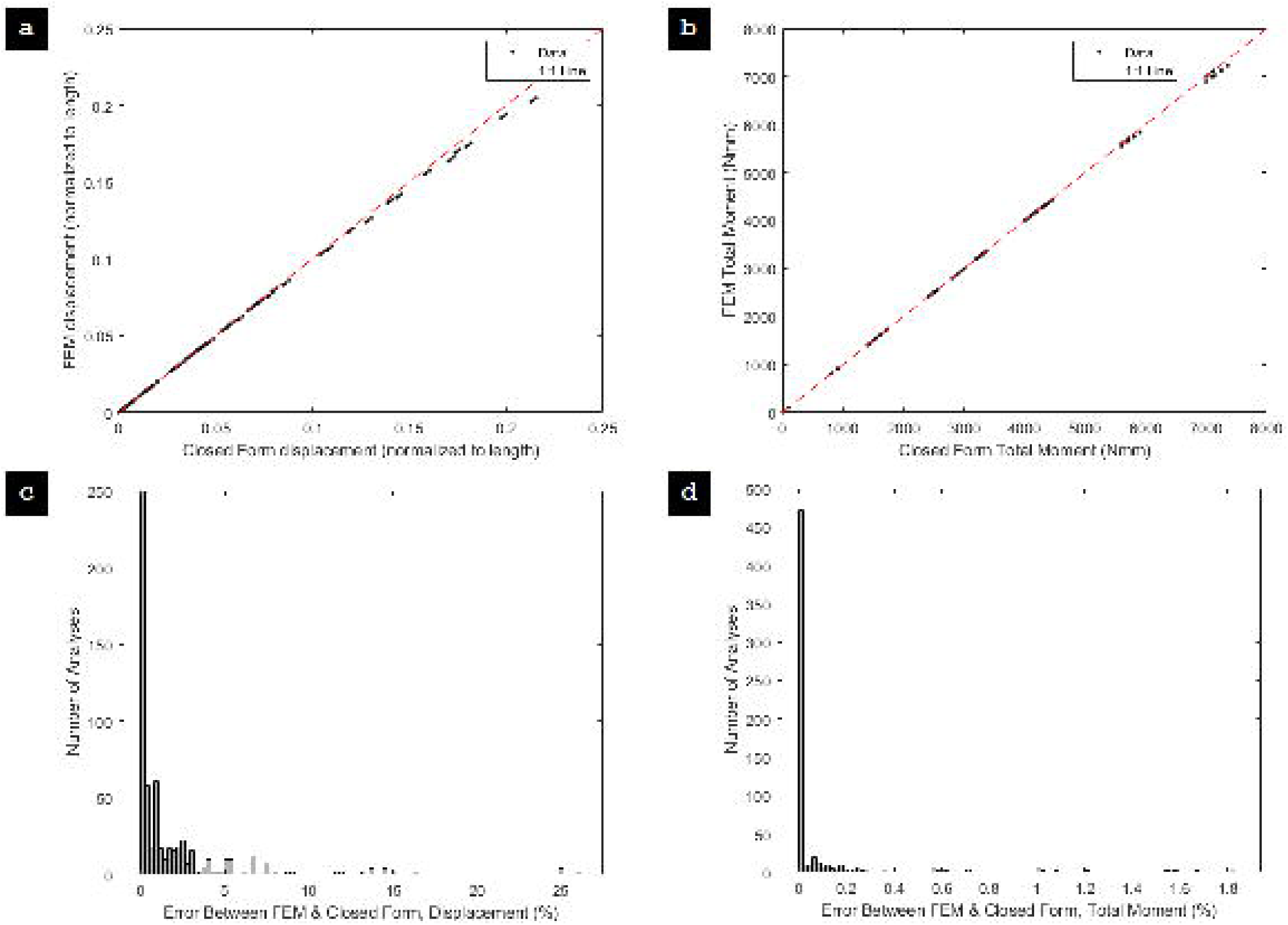
A comparison between the closed form solution and finite element models for the displacement at the top of the specimen (a) and the total moment at the base of the specimen (b), n= 768; A histogram of the error between the closed form solution and the finite element models for the displacement at the top of the specimen (c) and the total moment at the base of the specimen (d), n= 768.

An additional finite element model was created at the mean values in Table 1 to analyze the extent of applicability of the closed form solutions specifically to investigate extremely large deflections. Although this amount of deflection is not realistic in most plant stems, the model was analyzed to these extremely large displacements to demonstrate the nonlinear behavior and divergence of the closed form solutions beyond what would typically be seen in the field. Agreement between the finite element models and closed form solutions is strong at small displacements. At large displacements (greater than ∼45° at the tip), geometric nonlinearities that are not captured by the closed form engineering beam equations become more influential [4]. That is to say that the closed form solution is accurate so long as the linear closed form engineering beam equations are accurate. For more discussion on this topic, see the Limitations section. Figure 4 depicts the comparison between the finite element model with mean input parameters and the closed form solution. It should be noted that the model depicts a maximum horizontal displacement equal to the height of the stem.

**Figure 4:**
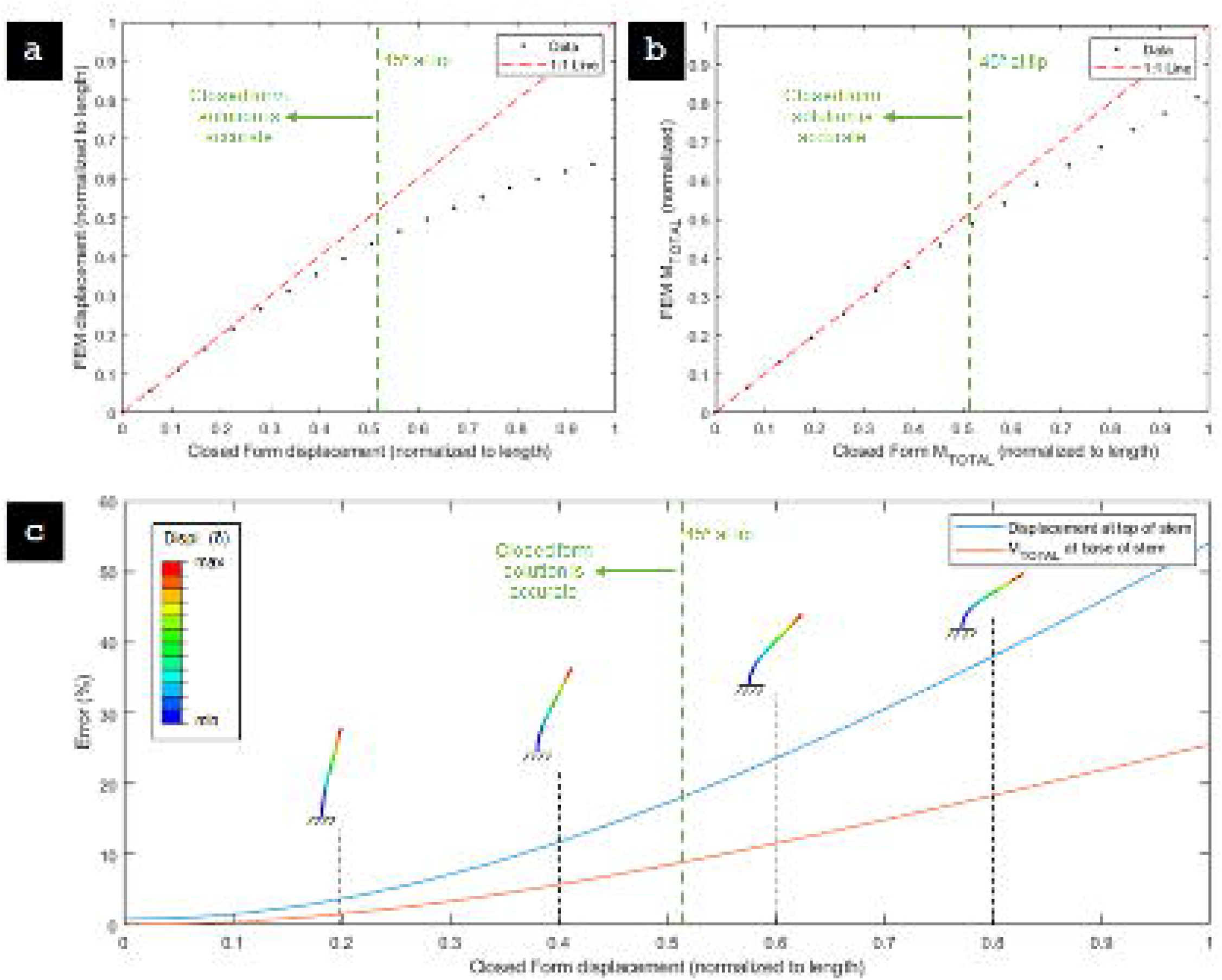
A comparison between the closed form solution and finite element model for displacements beyond loading that would typically be seen in the field. Plots depict the displacement (normalized to the height of the stem) at the top of the specimen (a) and the total moment (normalized to the maximum moment) at the base of the specimen (b); the error in the closed form displacement and total moment values for large displacements, with the finite element model displacements shown (c).

### A Computational Tool for Accounting for Weights

To make the closed form solutions more amenable to researchers without a structural engineering background (i.e., plant scientists, agronomists, and other end-users), an Excel (Microsoft Corporation, 2019) spreadsheet was developed, and is included in the Supplementary Data. The user simply inputs the flexural stiffness of the plant stem, the heights to each location of interest, the magnitude of externally applied forces and moments, and the weights at each location. Input values can be given for up to ten locations of interest along the length of the plant stem. The spreadsheet calculates the weight induced moments (M_int_), displacements, and the total induced moment (M_total_) at all locations. The spreadsheet makes the calculation both with and without self-loading considered. In addition, the error induced by ignoring the self-loading is calculated for the displacements and total induced moments. Additional instructions can be found in the spreadsheet. This tool can be used by researchers to determine the necessity of including self-loading in their studies. Figure 5 shows an example of the spreadsheet in which 3 externally applied forces, 2 internally applied moments, and 3 weights are considered.

**Figure 5:**
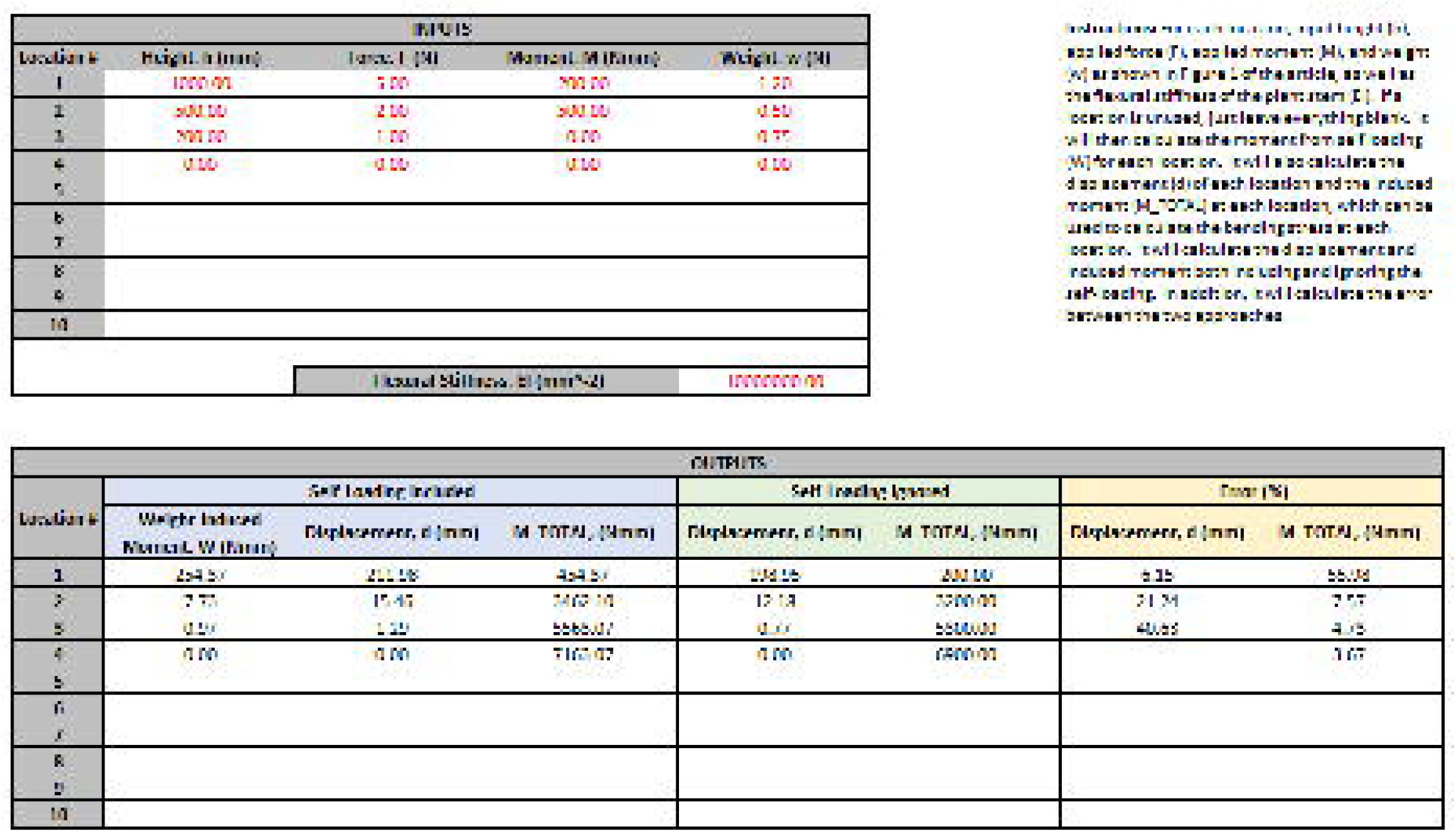
An example of the Excel spreadsheet in the Supplementary Data, showing loading at three locations, and calculating displacement and induced moments at four locations: the three loading locations and the base of the plant. Note that error in displacement is not calculated at the base, as displacement at the base is zero regardless of loading condition.

### Case Study: The Effect of Self-Loading on Wheat

As a demonstrative case-study, the effect of self-loading on wheat was analyzed. The flexural stiffness of wheat was assumed to be 0.027 Nm_2_ for this analysis [23]. The wheat biomass data collected as part of the current study (see methods section) was used to apply weights to the stem at every 10 cm along the length of the plant. A force of 1N was applied to the stem at the top of the stems (either 850mm or 950mm, depending on the sample). Figure 6 depicts the effect of self-loading as a percentage of the total displacement at the top of the stem and as a percentage of the total moment at the base of the stem. The contribution of self-loading was found to increase over the growing season, and then decrease during senescence. The contribution of self-loading was found to be significantly more for wet stems than dry stems, which is to be expected. As shown in Figure 6 self weight can account for up to 25% of the displacement of wheat stems and as much as 20% of the total moment.

**Figure 6:**
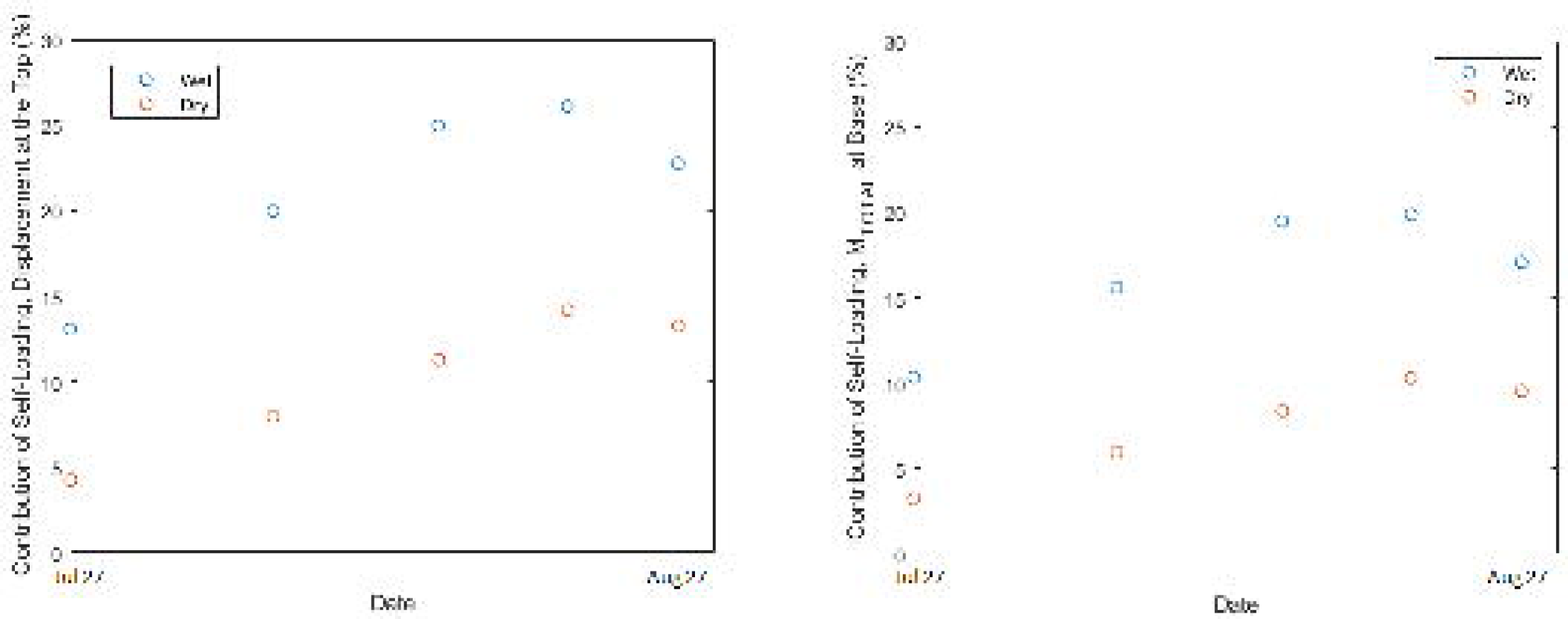
The contribution of self-loading for the displacement at the top of the stem (left) and the total moment at the base of the stem (right), for both wet and dry stems.

## Discussion

### The Influence of Self-Loading

The relative contribution of self-weight to the total bending moment can be conceptualized based on two key characteristics: (1) the displacement of the stalk, and (2) the ratio of weight to flexural stiffness (EI). This is because the moment induced by the weight is directly related to its offset (i.e., displacement of the stalk), as shown in Equation 6. The offsets (i.e., stem displacement) are in turn dependent on the flexural stiffness of the stalk. This means that a stalk with high flexural stiffness that requires a larger applied force to increase its displacement will be relatively less influenced by self weight. Conversely, a stalk with low flexural stiffness that requires a small amount of applied load to increase its displacement will be relatively more influenced by self-weight.

To aid researchers in determining if the influence of self weight is a significant factor in a given experiment the following examples are presented. These examples pertain to common mechanical phenotyping tests used to quantify flexural stiffness [24]. In addition, we present simple correction factors for the self weight-induced moments that are typically ignored in such experiments.

The first example loading configuration, which represents a typical flexural stiffness test for maize [6], applies a point load at the top of the specimen, while the stalk’s center of gravity is below the loading point. The second example loading configuration, which represents a typical flexural stiffness test for wheat [25, 26], applies a point load below the grain head but near the top of the specimen. Figure 7 displays the loading diagrams for these two test configurations.

**Figure 7:**
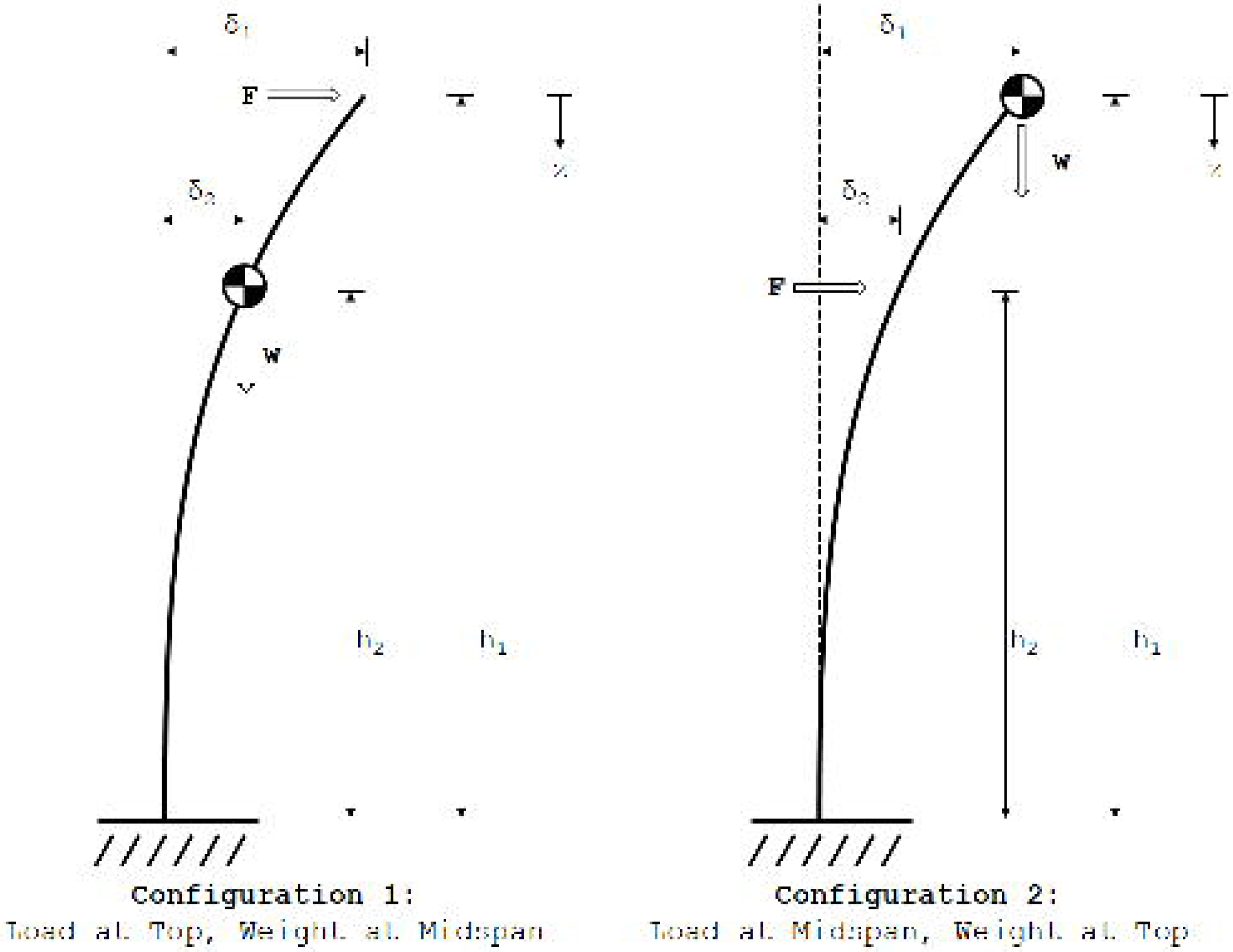
The loading diagrams for two common mechanical phenotyping test protocols used to determine flexural stiffness; a typical maize phenotyping protocol (left), and a typical wheat phenotyping protocol (right).

During these tests the applied load (F) and deflection of the stem at the point of loading (*δ*_1_) is typically recorded. Ignoring the weight of the stalk, the flexural stiffness of the stem (EI) can be calculated from the test data by rearranging the following equation to solve for EI:

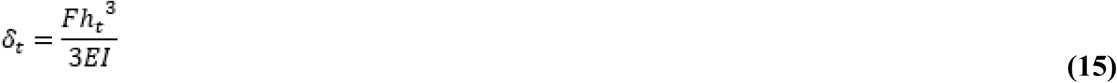

To account for the weight of the stalk when calculating the flexural stiffness of the stem, we must modify Equation 15 to include the stalk weight (w) as discussed in the methods section. For example:

#### Configuration 1: Load at Top, Weight at Midspan

First, solving Equation 11 for this loading configuration results in:

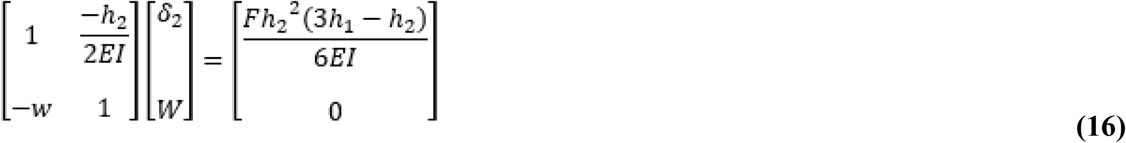

Where the two unknowns are the displacement at the weight (*δ*_2_) and the weight-induced moment (W). From this equation, the weight-induced moment can be calculated as:

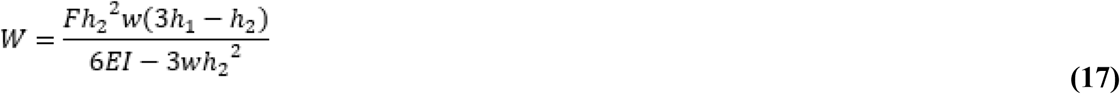

Finally, we can solve Equation 5 at the point of loading (*δ*_1_) to find a relationship between the test data and the flexural stiffness of the stem:

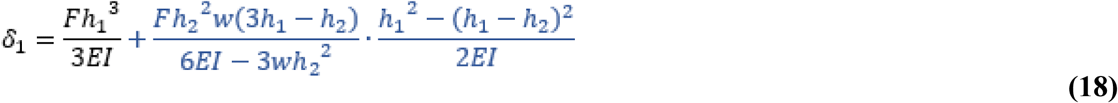

Where Equation 15 is shown in black, and the correction factor for the weight-induced moment is shown in blue.

#### Configuration 2: Load at Midspan, Weight at Top

As before, solving Equation 11 for this loading configuration at the weight location results in:

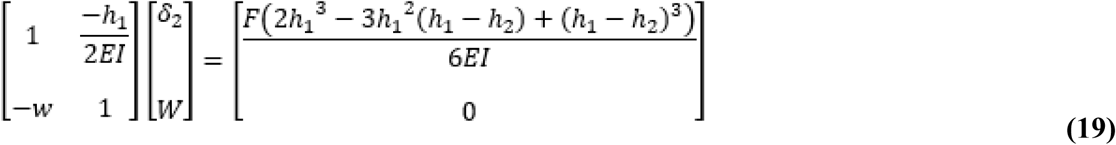

Solving for the weight-induced moment and solving for Equation 5 for the point of loading (*δ*_1_) to find a relationship between the test data and the flexural stiffness of the stem:

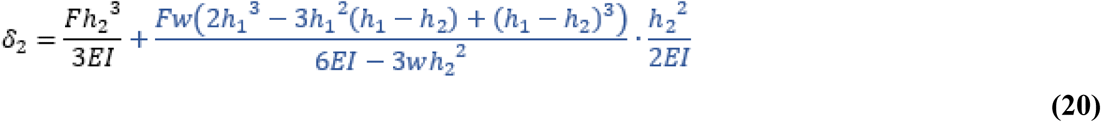

Where Equation 15 is shown in black, and the correction factor for the weight-induced moment is shown in blue.

It should be noted that because Equations 18 and 20 are simply Equation 15 with the addition of a correction factor that accounts for influence of the weight-induced bending moment, by comparing the results of Equation 15 with either Equation 18 or 20, the influence of the weight-induced bending moment can be calculated directly for each application. Thus, researchers can easily determine if the weight-induced bending moments are negligible for their testing purposes, or if they need to be incorporated into their biomechanical models.

### The Influence of Self-Loading for Various Plants

As a point of reference, five plants (maize, wheat, sweet sorghum, bamboo, and rice) were analyzed to determine the impact of the weight of the induced moments and displacements. The values shown in Table 2 represent typical values reported in the literature, but it should be noted that these are single data points only, and a significant amount of variation in heights, weights, and flexural stiffnesses is expected within a given plant. This information is presented here only as an accessible reference for researchers to develop an understanding of the types of plants that are more or less affected by self-loading.

**Table 2:** Self-loading related properties and the error of the induced moment at the base of the stalk if self-loading is ignored. The center of gravity of the plant is assumed to be halfway up the stem.

| Plant | Plant Height (mm) | Grain Height (mm) | Plant Weight (N) | Grain Weight (N) | Flexural Stiffness (Nm<sup>2</sup>) | Error of M<sub>TOTAL</sub> at base (%) | Error of displacement at top (%) | References |
| --- | --- | --- | --- | --- | --- | --- | --- | --- |
| Maize | 2250 | 1125 | 7.595 | 2.874 | 79.17 | < 1% | < 1% | [5,22,27–29] |
| Wheat | 638 | 638 | 0.016 | 0.021 | 0.027 | 12% | 16% | [23,30,31] |
| Sweet Sorghum | 2650 | 2650 | 9.64 | 0.346 | 137.1 | 1% | 1% | [32–34] |
| Bamboo | 10,774 | N/A | 138.02 | N/A | 229 | < 1% | < 1% | [35,36] |
| Rice | 969 | 969 | 0.0635 | 0.028 | 0.17 | 6% | 8% | [37,38] |

A key factor in determining the influence of self-loading is the ratio of the weight of a plant to its flexural stiffness. Although this parameter does not include all of the factors that influence self-loading, it can be used as a quick evaluation tool for researchers to determine the general amount of influence self-loading may have. Figure 8 depicts the influence of this ratio on the induced moment and displacement, with the plant varieties in Table 3 shown as data points. In general, it appears that self weight has a negligible effect on stiff and strong stems (i.e., bamboo and maize) but becomes more influential in smaller stems (i.e., rice, wheat).

**Figure 8:**
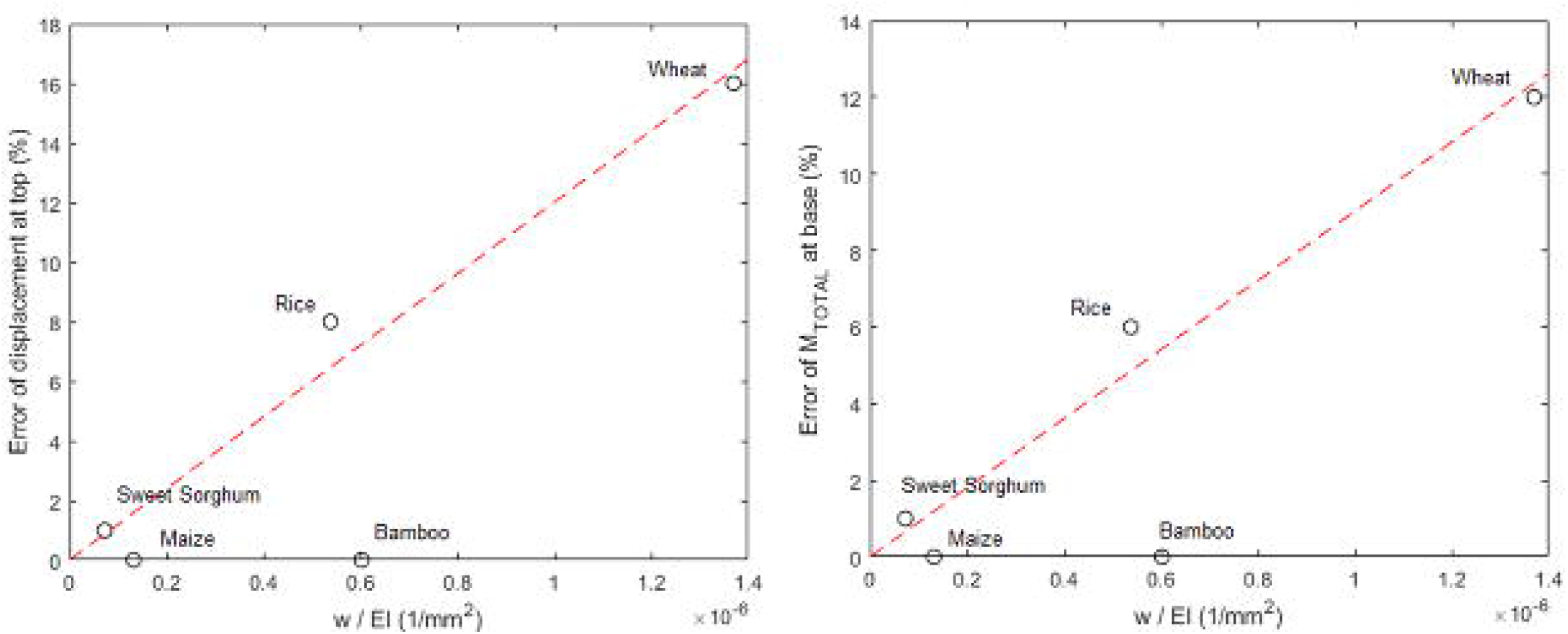
The error of displacement at the top of the stem (left) and M_TOTAL_ at the base of the stem (right), as a function of the ratio between the combined weight of the grain and plant and the flexural stiffness of the stem.

### Limitations

The primary limitation of the current study is that the stalk was assumed to be in-line with the assumptions made for pure bending, including maintaining a constant cross-section with homogeneous, isotropic, linear elastic material subjected to pure bending [4]. The inclusion of the changes in cross-sectional geometry along the lengths of the stalk [22], material heterogeneity and anisotropy, or non-linear material properties would likely change the behavior of the analytical system. Further discussion of the influence of such material assumptions on equations has been investigated in a previous study by the authors [4]. These assumptions, when combined with the assumption of a single cross-section along the entire length of the stalk, results in a single flexural stiffness parameter for the entire stalk. However, the flexural stiffness of plants changes constantly along the length of the stalk. This simplifying assumption was deliberately made to allow for an easily-used generalized equation. If researchers need to incorporate changes in flexural stiffness along the stalk, the approach presented in this study can be incorporated into a full Castigliano’s method beam approximation [17]. Additionally, the equations used in this study assume small strains and small displacements. As such, these equations carry the same limitations as standard engineering beam bending equations, and are not suitable to predict post-failure loading conditions or displacements. When post-buckling analyses are required, non-linear finite element modeling approaches are recommended.

Finally, Equations 1, 2, and 6 assume that the maximum moment induced by self-loading is applied to the entire length of the stem below the weight, which is not accurate, and is used as a simple estimation of the moment induced by self-loading. In reality, self-loading is not a constant moment along the length of the stalk, but instead is an axial compressive load that induces a moment that varies along the length of the stalk. However, modeling loading as an axial compressive load greatly increases the complexity of the equation, to the point that the matrix equations presented in this study would not be practical. Therefore, Equation 6 presents an upper-bound of the influence of self-loading by simply applying the maximum moment along the entire length of the stem. As shown in Figure 3 and Figure 4, this assumption is reasonable for the parameter space explored.

## Conclusions

The self-loading of plants plays a potentially critical role on the structural integrity of plant stems. To the best of the authors’ knowledge, this is the first study that presents a methodology for (1) incorporating the self-loading of plants with externally applied forces, (2) investigating the interconnected nature of the loading conditions placed on the plant, and (3) exploring the interconnected parameters that impact the influence of self-loading on the plant’s structural integrity. Two common phenotyping configurations were presented with simplified correction factors for researchers to use in their studies. A user-friendly Microsoft Excel framework was presented that allows researchers to quickly and easily determine the relevance of self-loading to their research experiments. A survey of five self-loaded plants was presented to give researchers an intuition on the level of influence self-loading plays on the structural integrity of various plants. It is the recommendation of the authors that self-loading be taken into account for plants such as wheat and rice that have a large ratio of weight to flexural stiffness.

## Conflicts of Interest

There are no conflicts to declare.

## Authors’ Contributions

All authors were fully involved in the study and preparation of the manuscript. The material within has not been and will not be submitted for publication elsewhere.

## Consent for Publication

Not applicable.

## Availability of Data and Materials

The datasets used and/or analysed during the current study are available from the corresponding author on reasonable request.

## Competing Interests

The authors declare that they have no competing interests.

## Acknowledgements

This work was funded in part by the National Science Foundation (Award #1826715) and by the United States Department of Agriculture - NIFA (#2016-67012-2381). Field data collection was completed by Undergraduate Research Assistants Matthew Kolbeck and Jonathan Fenske at the University of Saskatchewan’s Plant Phenotyping and Imaging Research Centre (P2IRC), which is supported through funding from the Canada First Research Excellence Fund (CFREF). Any opinions, findings, conclusions, or recommendations are those of the author(s) and do not necessarily reflect the view of the funding bodies.

